# Paralogous gene modules derived from ancient hybridization drive vesicle traffic evolution in yeast

**DOI:** 10.1101/2021.03.03.433305

**Authors:** Ramya Purkanti, Mukund Thattai

**Affiliations:** Simons Centre for the Study of Living Machines, National Centre for Biological Sciences, TIFR, Bangalore, India; Present address: Center for Integrative Genomics, Université de Lausanne, Lausanne, Switzerland

**Keywords:** evolutionary cell biology, organelles, gene duplication

## Abstract

Modules of interacting proteins regulate vesicle budding and fusion in eukaryotes. Distinct paralogous copies of these modules act at distinct sub-cellular locations. The processes by which such large gene modules are duplicated and retained remain unclear. Here we show that interspecies hybridization is a potent source of paralogous gene modules. We study the dynamics of paralog doublets derived from the 100-million-year-old hybridization event that gave rise to the whole genome duplication clade of budding yeast. We show that paralog doublets encoding vesicle traffic proteins are convergently retained across species. Vesicle coats and adaptors involved in secretory and early-endocytic pathways are retained as doublets, while tethers and other machinery involved in intra-Golgi traffic and later endocytic steps are reduced to singletons. These patterns reveal common selective pressures that have sculpted traffic pathways in diverse yeast species. They suggest that hybridization may have played a pivotal role in the expansion of the endomembrane system.

## INTRODUCTION

The eukaryotic vesicle traffic system is striking in its reliance on proteins encoded by large paralogous gene families. Distinct paralogs of Arf and Rab GTPases, vesicle coat proteins, cargo adaptors, and fusogenic SNAREs drive vesicle budding and fusion at distinct intracellular membranes [1]. The compositions of endomembrane organelles such as the endoplasmic reticulum, Golgi apparatus, and endosomes emerge dynamically, via the resulting loss and gain of vesicle cargo [2]. The organelle paralogy hypothesis posits that the generation of novel paralogs by gene duplication underlies the diversification of organelles over evolutionary time [3, 4]. This hypothesis is supported by phylogenetic evidence [5, 6] and biophysical modeling [2].

Vesicle traffic is regulated by membrane-specific modules of interacting proteins [7], so the emergence of new traffic pathways is likely to require the duplication of multiple genes within a module [2–4]. However, these genes are typically dispersed across the genome [8]. There are two ways an entire module can be duplicated: stepwise, via successive segmental duplications of its constituent genes [9]; or simultaneously, via a genome doubling event [10]. While some multi-protein complexes involved in vesicle traffic arose by stepwise segmental duplication [9], these represent only a small proportion of the full vesicle traffic apparatus. Little is known about the mode and temporal sequence of gene duplications by which the remaining paralogous modules arose. Here we ask whether genome doubling plays a significant role in the duplication and retention of vesicle traffic gene modules.

We examine the whole genome duplication (WGD) clade of budding yeast [11], which includes the well-studied vesicle traffic model *Saccharomyces cerevisiae*. This clade is the result of a 100-million-year-old hybridization event between two species belonging to distinct budding yeast lineages [12] (Fig. 1A). The offspring of such interspecies hybridizations are typically defective in meiosis, due to mismatches between homologous chromosomes. However, fertility is restored by a subsequent whole genome duplication event, which creates two identical copies of each chromosome that serve as homologous pairs during meiosis [13, 14] (Fig. 1B). The resulting allotetraploid cell, though functionally a diploid [15], has four copies of each gene: two distinct parental variants derived from hybridization, known as homeologs or paralogs; each present in two initially identical copies derived from WGD, which act as alleles (Fig. 1C).

**Fig. 1.**
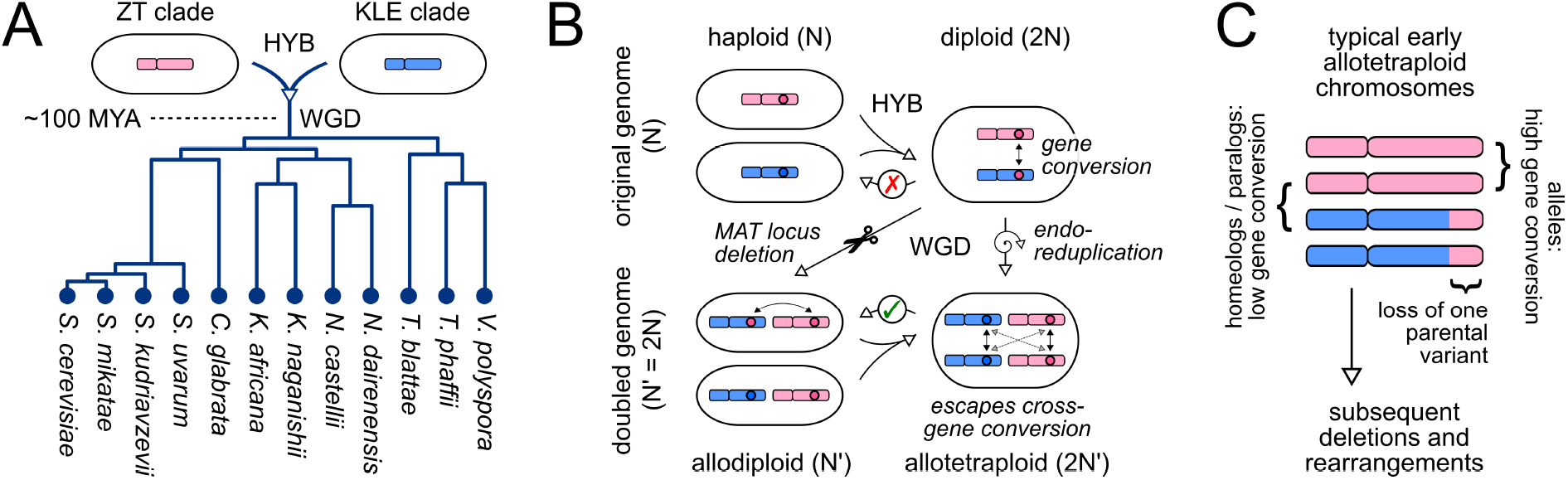
Generation of paralogs by interspecies hybridization. **(A)** The yeast whole genome duplication clade is descended from a hybridization between parents related to present-day members of the ZT and KLE clades. Phylogenetic branch lengths are from Ref. [11]. Branch lengths depicting interspecies hybridization (HYB) and subsequent whole genome duplication (WGD) are not to scale. **(B)** Routes to whole genome duplication in an interspecies hybrid. Pre-WGD hybrids are typically sterile (red cross) but genome doubling restores fertility (green tick). The genome doubling step can occur in one of two ways [13]: deletion of the MAT locus (diagonal arrow), which converts the pre-WGD diploid to a genome-doubled allodiploid (effectively a mating-competent haploid) [14, 26]; or endo-reduplication (vertical arrow), which converts the pre-WGD diploid to a genome-doubled allotetraploid (effectively a fertile diploid) [25]. In the hybrid diploid or allodiploid, gene conversion can replace one parental gene variant with another. In the allotetraploid, gene conversion occurs predominantly between the WGD-derived alleles rather than the hybridization-derived homeologs. **(C)** Typical organization of allotetraploid chromosomes soon after WGD. This schematic is based on the genome of the lager brewing yeast *S. pastorianus*, which is descended from a 500-year-old hybridization between *S. cerevisiae* and *S. eubayanus* [29, 30]. Most genes are present in two distinct homeologous variants. Loss of a parental variant is mostly restricted to contiguous stretches at chromosome ends, likely due to meiotic recombination in a pre-WGD hybrid diploid. In contrast, most allelic pairs are completely homozygous, either due to gene conversion or introgression.

The fates of these ancient paralogs, from their origins via hybridization to the present day, illuminate the evolutionary landscape of the vesicle traffic system. We find that the retention or loss of paralogs is highly correlated across members of the WGD clade, consistent with previous observations [16, 17] (Fig. 2). Among the paralogs retained as doublets, genes encoding vesicle traffic proteins are significantly enriched compared to the genomic background. There appears to be a consistent selective pressure to maintain specific paralogous modules as either doublets or singletons (Figs. 3,4). Vesicle coats and adaptors, and proteins that act along secretory and early-endocytic pathways, are typically retained as paralog doublets. In contrast, tethers and SNAREs, and proteins that act in intra-Golgi transport, late endocytic steps and vacuolar dynamics, are typically reduced to singletons. These patterns strongly support a scenario of convergent evolution in which members of paralogous gene modules are lost multiple independent times, but the probability of loss is low for modules involved in certain cellular activities. The implications are twofold: first, that hybridization is an important source of paralogous gene modules; second, that the availability of such modules directly impacts vesicle traffic. Though the yeast endomembrane system appears to be highly streamlined [18], our results reveal hidden layers of functional complexity sculpted by ongoing evolutionary dynamics.

**Fig. 2.**
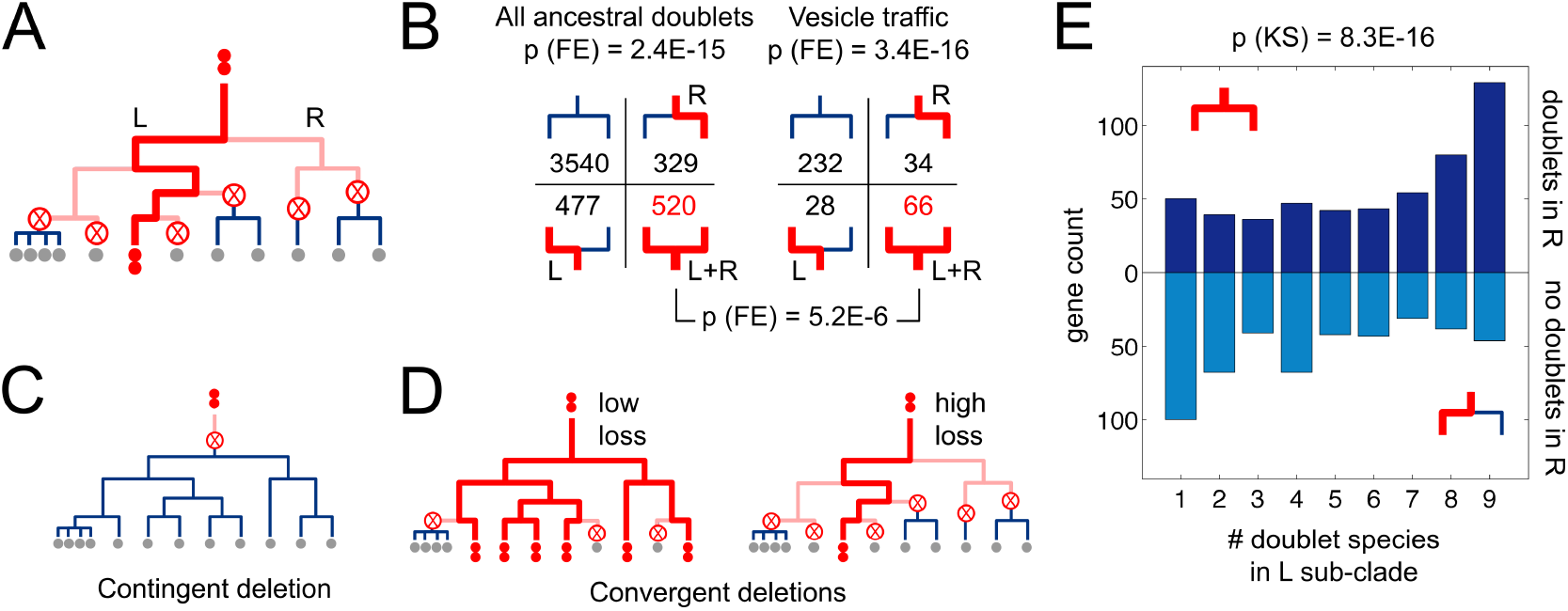
Patterns of paralog retention and loss. **(A)** Paralog doublets (double red dots) can be reduced to singletons (single gray dots) by loss of one gene copy (crossed circle). Bold red lines: lineages where presence of a doublet can be inferred, since a doublet is present in a descendant. Light pink lines: lineages where a doublet was present but this cannot be inferred, since the doublet is lost in all descendants. The cladogram is based on the phylogenetic tree from Fig. 1A, indicating the left (L) and right (R) sub-clades. **(B)** Each paralog can be placed into one of four groups depending on whether it is present as a doublet in any member of the L or R sub-clades. We show the total number of paralogs in each of these groups, for all 4866 ancestral doublets (left) and the 360 ancestral doublets encoding vesicle traffic proteins (right). P-value calculations are described in the main text and Methods. **(C,D)** Scenarios of doublet loss. **(C)** A single contingent deletion prior to the divergence of the clade. **(D)** Multiple convergent deletions on different branches, which can occur at low or high loss rates for different genes. **(E)** We test whether the presence or absence of doublets in the R sub-clade is predictive of the number of doublet species in the L sub-clade. We exclude cases in which there are zero doublet species in the L sub-clade, to ensure the doublet was not deleted prior to the divergence of the two sub-clades. The two histograms (dark blue, above x-axis, doublets in R sub-clade; light blue, below x-axis, no doublets in R sub-clade) are significantly different: a paralog present as a doublet in the R sub-clade is most likely to be a doublet in all species of the L sub-clade, whereas a paralog present as singletons in the R sub-clade is most likely to be a doublet in just one species of the L sub-clade. This supports the convergent scenario in Fig. 2D.

**Fig. 3.**
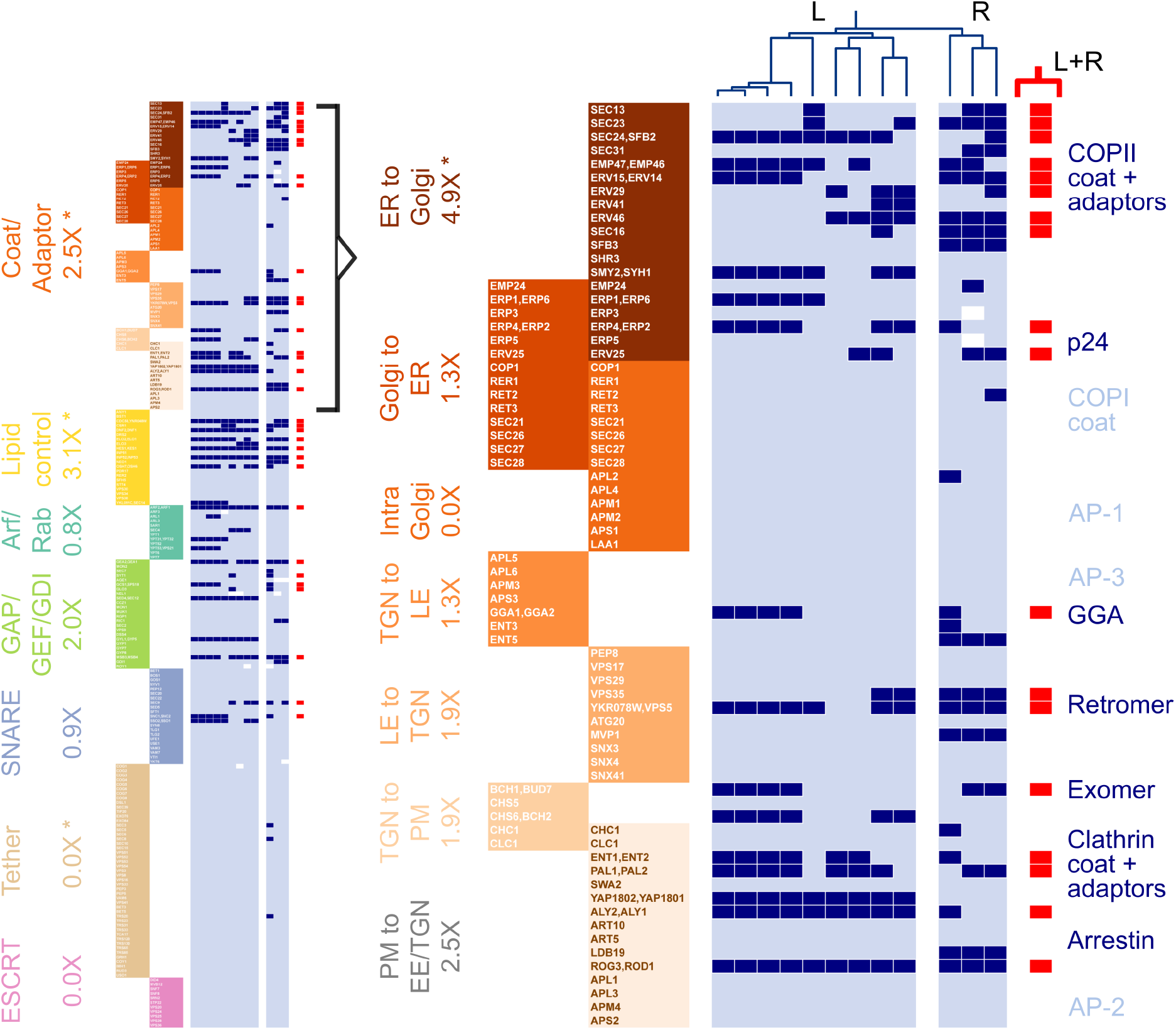
Paralogous vesicle traffic gene modules. We group 204 ancestral vesicle traffic genes into seven functional classes. We further sub-divide the Coat/Adaptor class into seven pathway-specific gene modules. ER: endoplasmic reticulum; PM: plasma membrane; EE/TGN: early endosome / trans-Golgi network; LE/PVE: late endosome / pre-vacuolar endosome. Note that some genes can act at multiple locations. Genes are labeled by the names of the corresponding *S. cerevisiae* homologs. We track whether each gene is present as a doublet (dark blue), a singleton (light blue), or has been completely lost (white) in each of the 12 species of the WGD clade. This information is represented as a matrix: rows correspond to genes, columns correspond to species. The L and R sub-clades are separated for visual clarity. The Coat/Adaptor portion of the matrix is shown expanded on the right. Paralogs present as doublets in both the L and R sub-clades are highlighted in red. Under each class or module description, we show the enrichment of L+R doublets compared to the background expectation. P-value calculations are described in the main text and Methods; * represents statistically significant enrichment or depletion.

**Fig. 4.**
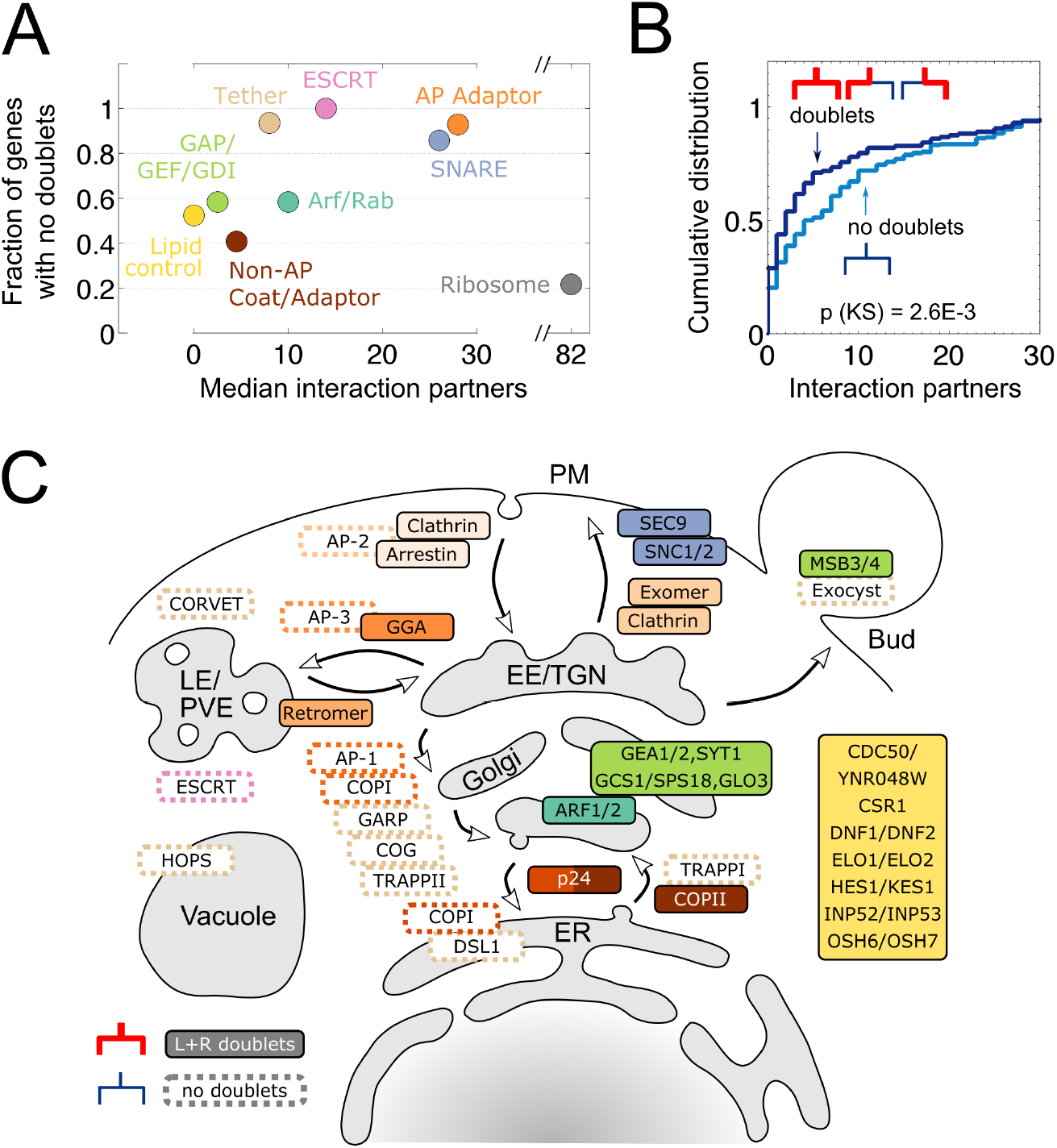
Landscape of vesicle traffic evolution in the WGD clade. **(A)** For each gene class from Fig. 3, we plot the median number of physical interaction partners for proteins encoded by the genes in that class (x-axis) against the fraction of genes in that class with no doublets in any species (y-axis). In this analysis, we have separated the AP Adaptor genes from the rest of Coat/Adaptor class. For comparison, we have plotted the corresponding values for ribosomal genes (gray dot; note the break in the x-axis). **(B)** We separate the 360 ancestral vesicle traffic doublets into two groups: those that are present as doublets in at least one species (dark blue) and those that are present as singletons in all species (light blue). Curves show the cumulative distribution of the number of physical interaction partners for proteins encoded by genes in each group. P-value calculations are described in the main text and Methods. **(C)** We show the site of action of proteins within the yeast vesicle traffic network. Filled boxes are proteins or complexes corresponding to genes present as L+R doublets. Dotted boxes show selected complexes for which the majority of proteins are present as singletons: the COPI coat and AP adaptors, tethers, and ESCRT.

## RESULTS

### Paralogs generated by hybridization escape gene conversion

Newly-generated paralog pairs, which typically have overlapping functions, may undergo neo- or sub-functionalization over time [19–21]. Before this happens, however, one copy may be transcriptionally silenced [22], deleted [23], or replaced by gene conversion [10]. Of these, gene conversion is the most rapid: it occurs during the repair of double-strand breaks, using an intact homologous sequence as a template [24].

For paralogs generated by genome doubling, the rate of gene conversion depends on whether the pre-WGD cell is a standard haploid or diploid, or an interspecies hybrid (Fig. 1B). Post-WGD cells are referred to as autopolyploids if they arose from a single species, or allopolyploids if they arose from an interspecies hybrid. Gene conversion is rapid in autodiploids [25] and autotetraploids [10, 26], since all paralogs are essentially identical. Gene conversion is also common in hybrid allodiploids [27, 28] since paralogs, though diverged, still represent the closest homologous pairs. In contrast to these cases, hybrid allotetraploids contain two classes of homologs: diverged homeologs inherited from distinct parental species, and near-identical allelic pairs arising from WGD (Fig. 1C). The availability of allelic pairs for double-strand break repair protects homeologous pairs from gene conversion [29, 30]. Though this increases the opportunities for neo- or sub-functionalization, there is no guarantee that both paralog copies will be retained.

### Most doublets in the WGD clade are retained as singletons

To track paralog retention, we obtained high-confidence paralog assignments in twelve members of the yeast WGD clade (Fig. 1A) from the Yeast Genome Order Browser [31] (ygob.ucd.ie; Methods). This dataset uses conserved gene order (synteny) to identify paralogs derived via the founding hybridization event of the WGD clade, distinguishing these from other homologous copies of genes within each genome which may have arisen via earlier or later duplication events.

For the remainder of our analysis we focus on the 4866 genes that comprise the ancestral paralog doublet set (Methods). Operationally, we define these as genes whose orthologs are found in the WGD clade, as well as in the ZT and KLE clades that represent the closest living relatives of the two species involved in the original hybridization [12] (Fig. 1A). This definition has good specificity (genes in this set are very likely to have been inherited as doublets in the original hybrid, one orthologous copy from each parent) and sensitivity (only 48/1374 = 3% of present-day doublets in any species fall outside this set). In *S. cerevisiae*, over 50% of present-day doublets map to pre-ZT or pre-KLE duplications (true paralogs inherited from both parents) and 25% of doublets map to duplications at the time of the WGD event (ohnologs derived from a single parent, likely the product of early gene conversions; Fig. 1C) [12].

Among the ancestral doublet set, 84% (4075/4866) are found in at least one copy across every WGD species. However, most of these have been reduced to singletons, with only ~10% persisting as doublets in a typical species [16]. This suggests that the 4866 genes in this set play important roles in single copy, but that the presence of a second paralogous copy is not typically advantageous, and may even be disadvantageous [32, 33].

### Vesicle traffic doublets are lost or retained convergently

The retention of some paralogs as doublets may be a transient stage in a slow process leading to eventual elimination of extra gene copies. Alternatively, there may be selective pressures favoring doublet maintenance (under neofunctionalization) or disfavoring doublet loss (under sub-functionalization). We can detect signals of such selection by comparing evolutionary trajectories across multiple species.

Since we are interested in long-term evolutionary patterns, we compared doublet occurrence in the two most distantly-related sub-clades of the WGD clade, which we refer to as the L and R sub-clades (Figs. 1A,2A; Methods). If a paralog is present as a doublet in any member of a sub-clade, we can infer it was present as a doublet at the root of that sub-clade (Fig. 2A, bold red lines). If a paralog is present as a singleton in every member of a sub-clade, we cannot draw any conclusions: one copy may have been deleted early, prior to the divergence of the sub-clade; or deleted later, independently in every member of the sub-clade (Fig. 2A, light pink lines).

Genes in the ancestral doublet set can be separated into four groups based on how they are retained in present-day species (Fig. 2B): those present as doublets in both sub-clades (L+R), those present as doublets in only one or the other sub-clade (L, R) or those present as singletons everywhere. Within the full gene set, doublet presence or absence is strongly correlated between the sub-clades (p = 2.4E-15, Fisher’s exact test; Fig. 2B, left). When we restrict the analysis to just those ancestral doublets involved in vesicle traffic [34] (pantherdb.org; Methods), the same correlation holds true (p = 3.4E-16, Fisher’s exact test; Fig. 2B, right). Even accounting for the genome-wide correlation of paralog doublets, vesicle traffic genes are significantly over-represented among the L+R set (66/360 = 0.18 compared to 520/4866 = 0.11; p = 5.2E-6, Fisher’s exact test). In summary, far more vesicle traffic paralogs than expected by chance are present either as doublets in both sub-clades, or as pure singletons in both sub-clades.

The observed doublet pattern could be contingent on history, the result of very early losses prior to the divergence of the sub-clades (Fig. 2C); or it could be convergent, the result of multiple later losses within each sub-clade (Fig. 2D). In a contingent scenario, the pattern arises purely due to shared ancestry, and is not connected to gene function. In a convergent scenario, the pattern arises because homologous genes have similar loss rates across different lineages. In this case, we expect a correlation in doublet loss between the sub-clades, conditioned on the doublet still being present in their last common ancestor. This conditioning is absent in the statistical tests done so far (Fig. 2B), but can be implemented as follows.

We pick a sub-clade, and only look at paralogs present as doublets in at least one member of that sub-clade; this enforces the condition that paralogs must be present as doublets in the last common ancestor of both sub-clades. For each paralog, we count the number of doublet species in this sub-clade. Finally, we split these paralogs into two groups, depending on whether or not they are present as doublets in some member of the other sub-clade. If the pattern is purely contingent, doublet species counts should be statistically indistinguishable between these two groups. Instead we find (Fig. 2E) that they are highly distinct (p = 8.3E-16 for L sub-clade counts, p = 4.2E-6 for R sub-clade counts, Kolmogorov-Smirnov test).

This analysis rules out pure contingency. It thus establishes that the observed convergent pattern arises because some types of genes are more likely to be retained as doublets, and others more likely to be reduced to singletons, with losses occurring independently in different species (Fig. 2D). This is consistent with prior observations that genes belonging to specific functional classes have been retained convergently across different species of the WGD clade [16, 17].

### Doublet retention varies across vesicle traffic gene modules

To explore the role of gene function in doublet retention or loss, we grouped genes into classes and modules. We define a vesicle traffic module as a set of genes whose protein products act collectively to carry out specific vesicle traffic functions at specific cellular locations [1, 7]. Within the ancestral doublet set, 360 are homologous to 420 present-day *S. cerevisiae* genes (300 singlets and 60 doublets) associated with the gene ontology term ‘vesicle-mediated transport’ [34] (pantherdb.org; Methods). Most of these doublets are true paralogs derived from hybridization [12], only 17 out of 60 trace their origins to the WGD event (the same proportion as among all present-day *S. cerevisiae* doublets).

We manually assigned these genes to seven functional classes (Fig. 3, left) based on annotations from the *Saccharomyces* Genome Database [35] (yeastgenome.org; Methods). We further sub-divided vesicle coats and adaptors into seven modules based on the traffic pathways where they are active [36] (Fig. 3, right). Out of 360 ancestral vesicle traffic doublets, we assigned 204 to classes and modules in this way. Of the remaining 156 genes, many play regulatory roles, and a few have not been precisely characterized.

Within each class or module, we asked how many paralogs were retained as doublets in both sub-clades (L+R doublets), and compared this to the expected number of doublets given the ~10% retention rate of the genomic background (Fig. 3). Among functional classes, coat/adaptor genes and lipid control genes were enriched for L+R doublets; and tethers and ESCRT genes had no L+R doublets. Among coat/adaptor modules, ER to Golgi traffic genes and PM to EE/TGN traffic genes were enriched for L+R doublets; and intra-Golgi traffic genes had no L+R doublets. Only four of these cases were statistically significant (Fisher’s exact test, Benjamini-Hochberg correction, FDR = 0.05; Methods): ER to Golgi traffic (4.9X enrichment, raw p = 6.6E-6); coats/adaptors (2.5X enrichment, raw p = 1.9E-4); lipid control genes (3.1X enrichment, raw p = 4.7E-3); and tethers (0.0X enrichment, raw p = 7.9E-3). The remaining classes were too small for the test to have statistical power.

### Modules with fewer physical interactions are slightly more likely to be retained as doublets

We next explored whether physical interactions influenced the retention of paralog doublets [33]. We imputed a physical interaction network among the proteins encoded by the ancestral doublet set, by using present-day interaction data for the corresponding proteins in *S. cerevisiae* [37] (string-db.org, only interactions based on curated databases or biochemical evidence were included; Methods). Across the gene classes defined previously (Fig. 3), we found that classes involving more physical interactions tended to have fewer genes retained as doublets (Fig. 4A). AP adaptors, SNAREs, ESCRT proteins and tethers, all of which have many interaction partners, were also the most likely to be reduced to singletons. Lipid control proteins and the non-AP coat/adaptor proteins, with fewer interaction partners, were most likely to be retained as doublets. However, the Arf/Rab class and their GAP/GEF/GDI regulatory proteins did not fit this pattern: both had equal rates of doublet retention, but the former had far more interactions than the latter. Moreover, for genes within each class there was no significant correlation between number of interaction partners and the number of paralog doublets.

To test this effect more rigorously, we separated all 360 ancestral vesicle traffic doublets into two groups: those reduced to pure singletons across all present-day species, and those present as doublets in at least one present-day species. Again, we found that the former group had more interaction partners than the latter group, and that this difference was statistically significant (p = 2.6E-3, Kolmogorov-Smirnov test; Fig. 4B). Statistically, this should be considered a weak effect since the inclusion of genes into these groups is not truly independent. Moreover, measured protein-protein interactions can be strongly influenced by abundance [38]. One cannot therefore rule out the possibility that doublet retention is driven by selective pressures causally independent of the number of interaction partners.

### Secretory and early endocytic pathways are enriched in paralogous modules

Finally, we considered the impact of paralogous gene modules in the context of the global yeast vesicle traffic system (Fig. 4C). Traffic pathways can be broadly classified into secretory components, from the ER via the Golgi and transGolgi network (TGN) to the plasma membrane (PM); and endocytic components, from the plasma membrane via early endosomes (EE) and late or pre-vacuolar endosomes (LE/PVE) to the vacuole. In *S. cerevisiae* the EE and TGN compartments appear to significantly overlap [18], serving as transit points during both secretion and endocytosis. Given these and other ambiguities about the sites of action of vesicle traffic proteins, it is difficult to formulate and statistically test hypotheses about whether paralogs involved in specific pathways are more likely to be retained as doublets. Nevertheless, the following patterns are suggestive of general principles.

Every step of secretion from the ER to the plasma membrane involves paralogous L+R doublets. At the ER to Golgi step, multiple components of the COPII coat and its adaptors [39], particularly the p24 complex [40], are L+R doublets. Within the Golgi, the master regulator ARF1/ARF2 is an L+R doublet, along with many Arf modulators involved in anterograde traffic such as GEA1/GEA2 [41]. At the TGN, cargo adaptors, including exomer which regulates TGN to PM export and GGA which regulates traffic to the PVE [36], are L+R doublets, along with components of the clathrin coat. At the plasma membrane, the v/t-SNARE complex comprising SNC1/SNC2 and SEC9, which drives fusion of secretory vesicles to the PM, are L+R doublets.

Aside from secretion, L+R doublets are involved in endocytic steps: early and intermediate clathrin coat proteins PAL1/PAL2 and ENT1/ENT2 are L+R doublets [42], along with components of the endocytic regulator arrestin. Two components of retromer, an adaptor which regulates cargo flow from the LE to the TGN [36], are L+R doublets. Another large class of L+R doublets are involved in regulating the synthesis and transport of lipids that play key roles in defining membrane identity. These include the lipid synthesis enzymes and regulators ELO1/ELO2, HES1/KES1, and OSH6/OSH7; lipid transport proteins CDC50, DNF1/DNF2, and CSR1; and signaling lipid regulators INP52/INP53.

In contrast to the cases discussed above, several modules involved in retrograde Golgi traffic and late endocytic steps have been completely reduced to singletons. These include: the COPI coat and all components of the AP adaptor complexes [36]; the entire set of coiled-coil and multi-subunit tethers [43]; and the ESCRT complex. COPI, AP-1, and the tethers GARP, COG, TRAPPII and DSL1, facilitate intra-Golgi cycling and Golgi-to-ER transport. The tethers CORVET and HOPS are involved in late endosomal and vacuolar dynamics, along with the ESCRT complex which remodels late endosomal membranes.

## DISCUSSION

### Hybridization as a source of paralogous modules

Polyploidy is a recurring theme in eukaryotic evolution, known to provide robustness against environmental perturbations, and to increase rates of niche adaptation [32]. These beneficial outcomes at ecological scales are the consequence of changes at the molecular scale. Here we have argued that polyploidy resulting from interspecies hybridization is an important source of paralogous gene modules. Interspecies hybrids among fungi are well documented [44, 45], and can spontaneously undergo WGD to restore fertility [13, 14]. These processes are not necessarily driven by selection for ploidy [32], but they do generate a large reservoir of paralogous gene modules as a byproduct.

Since paralogs that arise from hybridization are protected from gene conversion, they persist longer than initially identical gene copies arising from sporadic duplications [23] or non-hybrid whole genome duplications [26]. For paralogs that are parts of modules, a genome-scale duplication preserves the dosage balance of proteins and is less likely to perturb function, compared to a sporadic duplication [33]. These dynamics minimize the immediate deleterious effects of extra gene copies, promoting their maintenance in the short term. On longer timescales, however, extra gene copies tend to be lost unless they provide a specific selective advantage [32].

### Protein interactions influence the evolution of modules

The simultaneous duplication of many genes within a module sets the stage for neo- or sub-functionalization. However, it is not sufficient. Cross interactions between paralogous modules are common in newly formed yeast hybrids, even when parental species have diverged over 50 million years [46]. Tightly-interacting modules may be subject to dominant negative effects due to mutations in their paralogous partners, perhaps explaining why we see a weak association between interaction degree and doublet loss. This slightly deleterious effect can be offset by selection to maintain paralogs, as is observed for the yeast ribosome [47]: though this is the largest physically interacting complex in the cell, a majority of its genes are retained as doublets in most members of the WGD clade (Fig. 4A).

Cross interactions between paralogs are lost slowly over time, as proteins co-evolve and diverge from one another within the hybrid lineage [48]. Among modules reduced to singletons, one may expect the gene copies that are retained to be derived from the same parent, since these would have stronger internal interactions. In *S. cerevisiae*, nearly 70% of singletons are derived from the ZT clade, with the KLE variant being lost [12]. Among modules retained as doublets, the loss of cross interactions between distinct parental variants may actually be a prerequisite for gain of novel function [2].

### Selective advantage of paralogous vesicle traffic modules

A disproportionate number of paralogous gene modules have been retained in the yeast vesicle traffic system, 100 million years after they arose by hybridization. Though duplicate vesicle traffic modules have been described previously [49, 50], the fact many of these have a common origin has not been appreciated. It is only when these instances are identified and analyzed together that we truly see the scale and long-term impact of this ancient genome doubling event. Even more striking is the clear convergence of evolutionary trajectories across diverse yeast species, indicating common selective pressures operating on the vesicle traffic system over time and across ecological contexts. Yet the precise selective advantage of having paralogous vesicle traffic modules remains unclear. Paralog doublets may buffer the system against perturbations, or quantitatively increase the capacity of the secretory and endocytic routes. In support of this scenario, many L+R paralog doublets appear to have highly overlapping functions in *S. cerevisiae*. These include the COPII coat components EMP46/EMP47 [51], the TGN adaptors GGA1/GGA2 [52], the retromer components BCH1/BUD7 [53], and the clathrin adaptors ENT1/ENT2 [54]. However, not all paralogous pairs are redundant. The knockout phenotypes of certain paralogs are subtly different, as is the case for the arrestins ALY1/ALY2 [55] or the lipid control genes HES1/KES1 and OSH6/OSH7 [56]. Some paralog pairs are differentially expressed, with the more highly-expressed gene producing a more severe knockout phenotype. For example, the COPII components ERV14 and SEC24 are expressed at much higher levels than their respective paralogs ERV15 and SFB2 in *S. cerevisiae* [57, 58]. The Arf GEFs encoded by the doublet GEA1/GEA2 localize to different parts of the Golgi, with Gea1 in earlier compartments and Gea2 in later compartments [59]. The retromer component VPS5 has a paralogous copy in most WGD species whose function is completely unknown [60].

These examples suggest that the availability of paralogous modules may allow cells to tune their traffic pathways under different environmental conditions, or for different types of cargo. This is similar to what is observed for the yeast ribosome: its paralog doublets are differentially expressed under stress, and the resulting modulation of ribosomal composition confers a measurable fitness advantage [47].

### Early and late phases of endomembrane evolution

The last eukaryotic common ancestor (LECA) already had distinct versions of Arfs, Rabs, coats, and SNAREs which operated at distinct trafficking steps [3, 4]. These gene families arose very early in eukaryote evolution, over a billion years ago, and descendants of these proteins are found in all extant eukaryotic lineages. Compared to these early duplications, subsequent lineage-specific expansions appear fundamentally different in character. Though we find that many paralogs derived from hybridization have been retained within the vesicle traffic system, members of each paralog doublet invariably operate at the same trafficking step. Moreover, though most genes started off as paralog doublets following hybridization, many important classes of genes have been reduced to singletons convergently across multiple species. These results suggest that the architecture of vesicle traffic in present-day eukaryotes is tightly constrained, and that the hybridization route we have explored is distinct from pre-LECA duplication processes. It is likely the major vesicle traffic gene families were generated during an earlier, more dynamic and less constrained phase of eukaryotic evolution.

## ACKNOWLEDGEMENTS

We thank the participants of the 2015 KITP program on Evolutionary Cell Biology for discussions which nucleated this project. We particularly thank Andrew Murray for prompting us to consider the cell-biological context of genome doubling. We thank Jitu Mayor, Sunil Laxman, Benjamin Glick and Christian Landry for advice and feedback. We give special thanks to our families: without their patience and support through a year filled with challenges, this project could never have been completed.

## AUTHOR CONTRIBUTIONS

MT and RP designed the study. RP performed the bioinformatic analysis, MT performed the statistical analysis. MT wrote the paper.

## COMPETING INTERESTS

The authors declare that no competing interests exist.

## METHODS

### Ortholog assignments in pre-WGD and post-WGD species

We downloaded synteny-based ortholog assignments and paralog pair assignments from the Yeast Genome Order Browser [31] (YGOB Version 7; ygob.ucd.ie). This dataset covers twenty species: twelve within the yeast WGD clade, which we split into two sub-clades for further analysis (L subclade: *S. cerevisiae*, *S. mikatae*, *S. kudriavzevii*, *S. uvarum*, *C. glabrata*, *K. africana*, *K. naganishii*, *N. castellii*, *N. dairenensis*; R sub-clade: *T. blattae*, *T. phaffii*, *V. polyspora*); and eight pre-WGD species comprising the ZT clade (*Z. rouxii*, *T. delbrueckii*) and the KLE clade (*K. lactis*, *E. gossypii*, *E. cymbalariae*, *L. kluyveri*, *L. thermotolerans*, *L. waltii*). The WGD clade is descended from an interspecies hybridization between two species whose closest living relatives are inferred to belong to the ZT clade and the KLE clade, respectively [12]. A total of 14101 orthologs are present across all 20 species in the dataset. A subset of 11059 orthologs are found within the WGD clade.

### Defining the ancestral paralog doublet set

We are interested in orthologs that were present as paralog doublets immediately following the original interspecies hybridization. By definition, one copy of each such gene is inherited from each parent (Fig. 1B). However, we do not know the true genetic complement of the parental species, only that of their closest living relatives. Operationally, we define the set of ancestral doublets as the set of 4866 genes found across the ZT, KLE and WGD clades: these were definitely inherited from the parental species, and were very likely to have been inherited in two copies. We assigned a subset of *S. cerevisiae* doublets to pre-KLE or WGD duplication events based on Ref. [12]. The full list of ancestral doublets is provided in Supplementary Table S1.

### Annotation of vesicle traffic genes

We assigned genes to functional categories based on annotations of their *S. cerevisiae* homologs. 426 *S. cerevisiae* genes are annotated with the Gene Ontology term GO:0016192 ‘vesicle-mediated transport’ [34] (PANTHER Version 16.0; pantherdb.org). To these we added 17 genes whose paralogs were already part of the set. This resulted in 443 genes (323 singletons and 60 doublets in *S. cerevisiae*) of which 360 are present in the ancestral paralog doublet set (300 singletons and 60 doublets in *S. cerevisiae*). We used annotations from the *Saccharomyces* Genome Database [35] (yeastgenome.org) to assign 236 out of 443 genes (204 out of 360 ancestral vesicle traffic doublets) to seven functional classes: Coat/Adaptor; Lipid control; Arf/Rab; GAP/GEF/GDI; SNARE; Tether; and ESCRT. We sub-divided the Coat/Adaptor class into seven pathway-specific modules [36]: ER to Golgi (COPII, p24); Golgi to ER (COPI, p24); Intra Golgi (clathrin, AP-1, COPI); TGN to LE (AP-3, GGA, epsin); LE to TGN (retromer, nexin); TGN to PM (clathrin, exomer); and PM to EE/TGN (clathrin, AP-2, arrestin). Some Coat/Adaptor genes are active in more than one pathway. Annotations of vesicle traffic genes are provided in Supplementary Table S1.

### Statistical tests

We performed all enrichment analyses (Figs. 2B, 3) using the two-tailed Fisher’s exact test on 2×2 contingency tables. When testing for enrichment among the 14 vesicle traffic gene classes/modules (Fig. 3), we additionally applied the Benjamini-Hochberg correction for multiple hypothesis testing with a false discovery rate *α* = 0.05, to determine the significance threshold. For comparing between distributions (Figs. 2E, 4B) we used the Kolmogorov-Smirnov test. Raw data and p-values for each test are provided in Supplementary Table S2.

### Paralog protein interaction network

The 360 ancestral vesicle traffic genes correspond to 420 *S. cerevisiae* genes (300 singletons and 60 doublets). For these genes we obtained the protein-protein interaction network from the STRING database [37] (string-db.org), filtering for the physical sub-network at medium confidence, with experiments and databases as interaction data sources. For genes present as doublets, we assumed an interaction between a pair of ancestral genes if there was an interaction between any of their paralogs, as would be expected based on a subfunctionalization scenario. For comparison, we repeated this analysis for 83 ancestral ribosomal genes, corresponding to 138 *S. cerevisiae* genes (28 singletons and 55 doublets).

## SUPPLEMENTARY INFORMATION

Raw data are provided in Supplementary Tables S1 and S2.

## Notes

### Competing Interest Statement

The authors have declared no competing interest.

